# OligoFormer: an accurate and robust prediction method for siRNA design

**DOI:** 10.1101/2024.02.02.578533

**Authors:** Yilan Bai, Haochen Zhong, Taiwei Wang, Zhi John Lu

## Abstract

**Motivation:** RNA interference(RNAi) has become a widely used experimental approach for post-transcriptional regulation and is increasingly showing its potential as future targeted drugs. However, the prediction of highly efficient siRNAs(small interfering RNA) is still hindered by dataset biases, the inadequacy of prediction methods, and the presence of off-target effects. To overcome these limitations, we propose an accurate and robust prediction method, OligoFormer, for siRNA design.

**Results:** OligoFormer comprises three different modules including thermodynamic calculation, RNA-FM module, and Oligo encoder. Oligo encoder is the core module based on the transformer encoder. Taking siRNA and mRNA sequences as input, OligoFormer can obtain thermodynamic parameters, RNA-FM embedding, and Oligo embedding through these three modules, respectively. We carefully benchmarked OligoFormer against 5 comparable methods on siRNA efficacy datasets. OligoFormer outperforms all the other methods, with an average improvement of 9% in AUC and 10.7% in F1 score in our inter-dataset validation. We also provide a comprehensive pipeline with prediction of siRNA efficacy and off-target effects using PITA score and TargetScan score. The ablation study shows RNA-FM module and thermodynamic parameters improved the performance and accelerated convergence of OligoFormer. The saliency map by gradient backpropagation shows certain base preferences in initial and terminal region of siRNAs.

## 1 Introduction

RNAi makes siRNAs promising therapeutic drugs due to their potential to silence disease-related genes in a sequence-specific manner[1]. siRNAs are introduced into the RNA-induced silencing complex (RISC), which comprises several distinct proteins, including Argonaute-2 (Ago-2), Dicer, and the transactivation response element RNA-binding protein (TRBP)[2]. After the activation of siRNAs through removal of their sense strands, the resulting antisense strands direct the RISC to bind with the target mRNA, where Ago-2 facilitates cleavage[3]. Upon cleaving the target mRNA, the siRNA-loaded RISC can dissociate and engage with another mRNA molecule. Consequently, minimal concentrations of siRNAs effectively induce gene knockdown. Notably, siRNAs exert their influence post-transcriptionally at the mRNA level, offering a distinct advantage over post-translational protein-focused approaches. This feature allows for the targeting of “undruggable genes” for which inhibitors are unavailable or challenging to develop, significantly expanding the range of potential therapeutic targets beyond traditionally druggable proteins[4]. As of 2022, 10 RNAi drugs have been approved by the FDA or entered late-phase 3 clinical trials[5], indicating the great potential of RNAi drugs in the future.

However, obtaining highly efficient siRNAs is a challenging task, requiring specific characteristics such as the absence of innate immune system activation, high efficacy in cutting specific targets, and minimal off-target or toxic effects[4]. Hence, the design of effective siRNAs is crucial for the success of RNAi therapeutics, leading to the development of various computational tools to aid in this process. While many tools, such as OligoWalk[6], siRNAPred[7], and i-score[8], have been tested and demonstrated effectiveness, they still exhibit some shortcomings. These can be broadly discussed in terms of dataset biases, prediction methods, and off-target effects.

Dataset biases may affect the generalization and validity of siRNA design models. Huesken et al.[9] collected 2,431 siRNAs targeting 34 mRNAs along with their inhibition values, making a significant contribution to the enrichment of siRNA datasets. Some models, such as LASSO-based regression model by Vert et al.[10], were trained primarily on this dataset.

While they achieved rather good results, they lacked validation on other datasets, raising concerns about their generalization ability. Another important consideration is the significant variation in experimental conditions and data quality among different datasets. For example, Huesken et al. utilized a high-throughput fluorescent reporter gene system, while Hsieh et al. employed quantitative real-time PCR for measurement[9, 10, 11]. Normalizing data across diverse datasets and integrating them appropriately pose significant challenges.

The effectiveness of the final result is influenced by the characteristics of the prediction methods or models. Early researchers endeavored to employ machine learning methods for the analysis of siRNA efficacy. Huesken et al. used the Stuttgart Neural Net Simulator to train an algorithm named BIOPREDsi[9]. Ichihara et al. developed a simple linear regression model, i-Score, to predict active siRNAs while considered only nucleotide preferences at each position[8]. Lu et al. constructed a support vector machine (SVM) that selects functional siRNAs based on both thermodynamic and sequence features[12, 13]. While these methods achieved good results at the time, the structures of these models were relatively simple, and they could not extract some hidden features well. With the rise and proven capabilities of deep learning, researchers started applying deep learning models to biological problems, aiming to capture deep-dimensional features for analysis. Han et al. utilized convolution kernels as motif detectors to extract siRNA sequence features. After combining thermodynamic properties with a pooling layer, they introduced a deep neural network (DNN) to generate feature representations and output efficacy through a logistic regression function[14]. The performance of their proposed method was improved on the same dataset of Huesken et al. Apart from this, Massimo et al. proposed a graph neural networks (GNN) approach to simulate the siRNA-mRNA interaction networks. siRNAs and mRNAs are encoded with 3-mer and 4-mer fragments respectively as two types of nodes whereas 22 thermodynamic parameters calculated from RNAUp web server and Gibbs energy serve as the third node[15]. They took the prediction of siRNA efficacy to a new level. However, due to the limited complexity of these models, they still struggled to capture the binding properties between the long context of mRNAs and siRNAs. In recent years, transformer-based models have shown remarkable success in various natural language processing and genomics tasks compared to earlier deep learning models[16]. siRNA sequences share similarities with sentences in language tasks, making transformers a probable choice. Furthermore, pre-trained large models are starting to demonstrate their powerful capabilities in downstream migration tasks. RNA-FM is the first foundation model for the community to accommodate all non-coding RNA sequences[17]. These pre-trained RNA models may enrich the feature representation of siRNA-mRNA interactions.

Off-target prediction is quite vital for the practical application of siRNAs in treatment. However, many existing prediction models only focus on the effectiveness of binding predicted siRNAs to target mRNA to produce inhibitory effects, overlooking the potential off-target effects caused by the siRNA sequence itself. The off-target effects associated with siRNAs delivery can be divided into three broad categories: miRNA-like off-target, immune stimulation and saturation of the RNAi mechanism[18]. siRNAs, with seed region complementarity acting like miRNAs, may down-regulate a large number of transcripts[19]. While chemical modification of the seed region can reduce the effects[20], a comprehensive transcriptome comparison remains necessary. For predicting the miRNA-like off-target effects, well-established miRNA target prediction tools can be applied in this field[21]. Immune stimulation mainly results from specific motifs in the siRNA strand, such as UGUGU, GUCCUUCA, and CUGAAUU[22]. Regarding RNAi mechanism saturation, no strategies have been identified to mitigate this effect[18].

Here we propose a novel method for predicting siRNA efficacy and off-target effects, named OligoFormer. This method consists of two components: a transformer-based model to capture deep hidden sequence features and learn complex patterns of siRNA-mRNA interactions for siRNA efficacy prediction, and an overall pipeline to select the best siRNA predicted by our model, taking into account various off-target effects.

## 2 MATERIALS AND METHODS

### 2.1 Dataset collection and preprocessing

#### 2.1.1 Dataset collection

We strategically aggregated a diverse array of datasets to facilitate accuracy and robustness of our model. We collected 9 datasets comprising a total of 3,714 siRNAs and 75 mRNAs from previous studies, including those by Huesken[9], Takayuki[23], Amarzguioui[24], Haborth[25], Hsieh[11], Khvorova, Reynolds[26], Vickers[27], and Ui-Tei[28](Table 1). We categorized the datasets into three sets: the Huesken dataset, the Takayuki dataset, and merged the remaining datasets as Mixset (Table 1S). The inhibition efficacy or activity of siRNAs in all datasets were normalized into inhibition efficiency ranging from 0 to 100%, and 70% of the maximum inhibition was used as the threshold to classify positive and negative siRNAs.

**Table 1.** The datasets of siRNAs.

| Datasets | siRNAs number | mRNAs number | Cell line |
| --- | --- | --- | --- |
| Huesken | 2431 | 34 | H1299 |
| Takayuki | 702 | 1 | HeLa |
| Amarzguioui | 46 | 4 | HaCaT |
| Haborth | 44 | 1 | HeLa |
| Hsieh | 108 | 22 | HEK293T |
| Khvorova | 14 | 1 | HEK293 |
| Reynolds | 240 | 7 | HEK293 |
| Vickers | 76 | 2 | T24 |
| Ui-Tei | 53 | 3 | HeLa |

#### 2.1.2 Sequence normalization

To facilitate model training, it is essential to ensure consistent lengths for both siRNAs and mRNAs across different datasets. Specifically, we uniformly truncated siRNA sequences to 19 nucleotides. In most datasets, siRNAs were naturally 19 nucleotides long, while in the Huesken dataset, they were 21 nucleotides long and the 2 nucleotides at positions 20-21 in the 3’-overhang were truncated to ensure uniformity. For mRNA sequences, we extended 19 nucleotides upstream and downstream to reach a total length of 57 nucleotides, based on the complementary siRNA sequence. Extensions of upstream and downstream parts less than 19 nucleotides are padded with the X nucleotide. This standardization method enables consistent processing and feature extraction across all datasets.

#### 2.1.3 RNA-FM Model Embedding

In this study, we utilized RNA language model RNA-FM to generate embeddings for both siRNAs and mRNAs, serving as additional features. RNA-FM can extract various structural and compositional features of RNAs, capturing nucleotide information from both the primary sequence and secondary structure. The embeddings generated by RNA-FM were used as additional high-dimensional input features for OligoFormer, enhancing the model’s ability to learn relationships between siRNAs and mRNAs. The shape of the embeddings is determined by the length of the RNA sequence multiplied by 640. Therefore, the shape of the siRNA embedding is 19×640, and the shape of the mRNA embedding is 57×640. All RNAs in our datasets were embedded and pre-saved in the *npy* format using RNA-FM for subsequent model training.

#### 2.1.4 Thermodynamic Parameters

In addition to standard sequence embeddings, OligoFormer integrates RNA thermodynamic parameters as crucial input features. These features (Fig.1(a);Table 2S) were computed for each siRNA sequence, providing valuable insights into the thermodynamic stability and binding affinity of siRNA-mRNA interactions. The nucleotide content provides information about the stability of the interaction, indicating the conditions under which the siRNA effectively binds to the mRNA. Furthermore, Δ*G* quantifies the free energy change associated with the formation of the siRNA itself and the duplex, providing a measure of the thermodynamic favorability of the binding. Lower values of Δ*G* suggest more energetically favorable interactions. These thermodynamic parameters represent some intrinsic properties of RNA and provide a comprehensive foundation for siRNA efficacy prediction. Integrating them into the model’s input features allows OligoFormer to gain a better understanding of the energetics governing siRNA functionality.

**Fig. 1.**
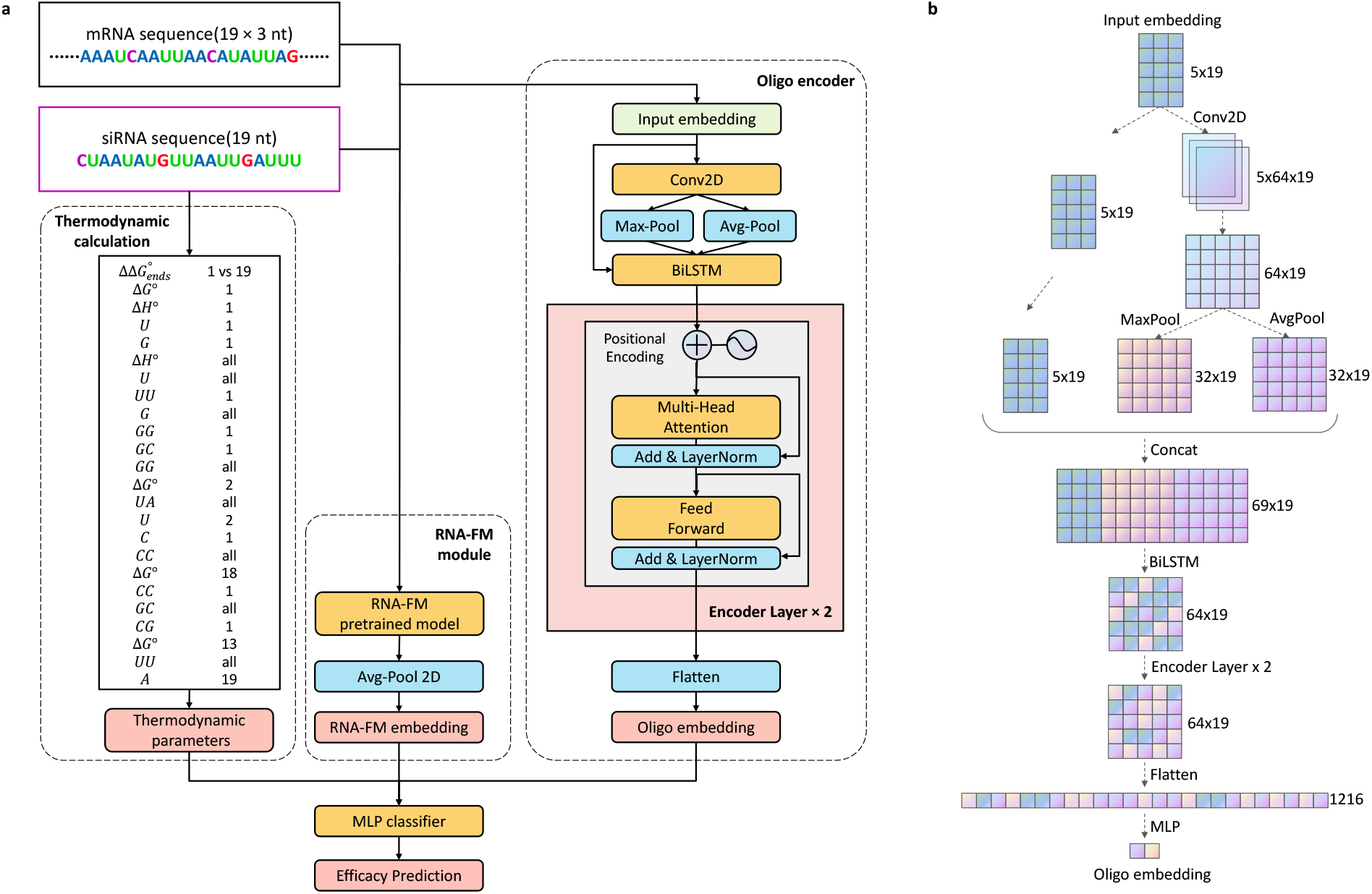
The overview of OligoFormer. (a)The architecture of OligoFormer. All features including thermodynamic parameters, RNA-FM embedding, and Oligo embedding, are first calculated from siRNA and mRNA sequences through different modules and then concatenated together into MLP classifier to obtain the prediction of siRNAs. The Oligo encoder comprises a 2D convolution layer, a max pooing layer, an average pooling layer, a bidirectional LSTM, two-layer multi-head transformer encoder, and a flatten layer to get Oligo embedding. (b) The computational graph illustrates the detailed structure of Oligo encoder and the shape of RNA embedding at each layer of Oligo encoder, using 64 as the example dimension of the hidden layer.

### 2.2 The architecture of OligoFormer

As illustrated in Fig.1(a), OligoFormer takes siRNA and mRNA sequences as input and comprises three different modules including thermodynamic calculation, RNA-FM module, and Oligo encoder. Thermodynamic parameters, RNA-FM embeddings and Oligo embeddings will be calculated from individual processing through these modules and OligoFormer employs a late fusion mechanism to combine diverse features by concatenated into a Multilayer Perceptron(MLP) classifier. This architecture allows OligoFormer to capture both individual features and their combined impact on siRNA efficacy. A pivotal module of OligoFormer is Oligo encoder, which is responsible for distilling complex information from the input features and generating a comprehensive Oligo embedding. The detailed structure of the Oligo encoder consists of a sequence of layers tailored for different aspects of feature extraction. Starting with a 2D convolutional layer, the encoder captures spatial dependencies within the input features, which is followed by a max pooling layer and an average pooling layer, extracting features while retaining the most relevant information. The Bidirectional Long Short-Term Memory(BiLSTM) layer introduces temporal dynamics and captures sequential dependencies. Two-layer multi-head transformer encoder further enhances the model’s ability to capture complex patterns by leveraging attention mechanisms. Lastly, a flatten layer consolidates the extracted features to obtain a comprehensive RNA embedding. To visually illustrate the embedding flow, Fig.1(b) presents a computational graph detailing the shape of the RNA embedding at each layer of the Oligo encoder. With a dimension of 64 for the hidden layers, the graph illustrates how the features change from input embedding to Oligo embedding.

### 2.3 Cross-Validation Methods

We compared OligoFormer with previous open source methods, including OligoWalk, siRNAPred, i-score, s-Biopredsi, and DSIR on both intra-dataset and inter-dataset validation. Other softwares of commercial companies for siRNA design, such as siRNA wizard[29], siDesign Center[30] and siRNA Target Finder[31], failed to open source, so their relative effectiveness cannot be evaluated here.

For intra-dataset validation, we employed a 5-fold cross-validation strategy on the Huesken, Mixset, and Takayuki datasets to mitigate biases and ensure robustness. Each dataset was split into 5 subsets, with each subset serving as a validation set exactly once to evaluate models’ performance across diverse data splits. OligoFormer and the other 5 methods were employed using this validation scheme to assess their performance on individual datasets, which provided a basic standard to measure the accuracy and robustness of these models.

For inter-dataset validation, we employed it to assess the model’s generalization and robustness across different datasets. We trained models on the Huesken dataset and evaluated their performance on the Mixset dataset. The ability of models to provide accurate predictions across diverse datasets is crucial for their applicability in real-world scenarios with varying experimental conditions.

## 3 RESULTS

### 3.1 Performance on intra-dataset validation

We compared OligoFormer with other siRNA design models, such as OligoWalk, siRNAPred, i-score, s-Biopredsi and DSIR. The performance of these models was evaluated on three datasets, Huesken, Mixset, and Takayuki, using 5-fold cross-validation. AUC (area under curve) and F1 score were used as metrics to measure model performance. As Fig.2 shows, OligoFormer outperforms other methods on all three datasets, achieving an average AUC of 0.8619 and an average F1 score of 0.7584 on Huesken, an average AUC of 0.8444 and an average F1 score of 0.7636 on Mixset, and an average AUC of 0.8628 and an average F1 score of 0.577 on Takayuki. OligoFormer stands out among other methods on intra-dataset validation, demonstrating its accuracy in learning and understanding one certain dataset.

**Fig. 2.**
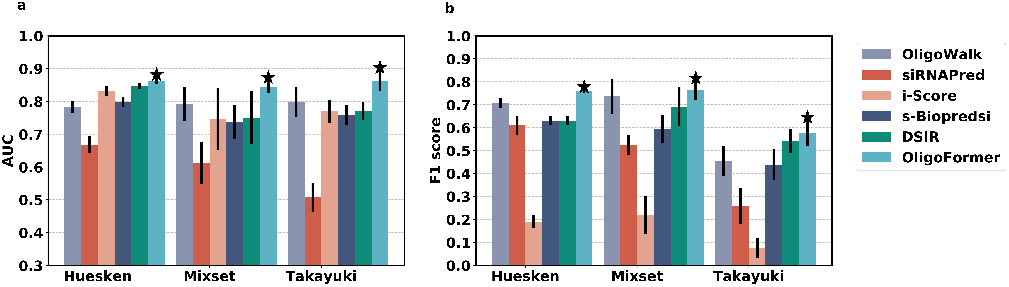
The performance on intra-dataset validation. (a) The AUC values of OligoFormer and the other 5 methods on Huesken, Mixset, and Takayuki datasets in 5-fold validation. The black vertical line represents standard error bar, and the star above it indicates the best AUC. (b) The F1 scores of OligoFormer and the other 5 methods on Huesken, Mixset, and Takayuki datasets in 5-fold validation.

### 3.2 Performance on inter-dataset validation

In addition to the ability to understand one certain dataset, the model’s predictive performance across datasets is even more crucial. In our inter-dataset comparison, OligoFormer was trained on the Huesken dataset and validated on the Mixset and Takayuki datasets, alongside OligoWalk, siRNAPred, i-score, s-Biopredsi, and DSIR. AUC and F1 score were used to evaluate the performance of each model across different datasets. As illustrated in Table 2, OligoFormer outperforms the other models on all valid datasets, achieving an average AUC of 0.8153 and an average F1 score of 0.7689 on the Mixset dataset. Therefore, OligoFormer is more robust for varying sequence characteristics, experimental conditions, and assay methodologies than other methods.

**Table 2.** Performance on inter-dataset validation.

| Methods | AUC | F1 score |
| --- | --- | --- |
| OligoWalk | 0.708 | 0.6258 |
| siRNAPred | 0.5782 | 0.5245 |
| i-Score | 0.7462 | 0.2197 |
| s-BioPredsi | 0.7374 | 0.5936 |
| DSIR | 0.7483 | 0.6945 |
| OligoFormer | **0.8153** | **0.7689** |

### 3.3 Ablation study

Input features and modules of OligoFormer were analyzed through an ablation study based on inter-dataset validation. First, the essentiality of 8 different features, including siRNA, mRNA, RNA-FM, and thermodynamic parameters, along with their combinations, was evaluated using AUC and F1 score. As Fig.3(a) shows, the combination of all input features achieved the best performance, followed by the combination of siRNA, mRNA, and RNA-FM model. Removing any module will cause the model to become less accurate. RNA-FM module improved the model’s performance because it is able to extract deep-dimensional features of siRNA-mRNA interactions compared to traditional models. The incorporation of TD(thermodynamic parameters) enriches the features used for prediction and allows OligoFormer to capture a broader spectrum of information, enabling a more accurate evaluation of siRNA efficacy. Using a 19-nucleotide sliding window to scan the entire target mRNA sequence is another distinctive feature of OligoFormer, ensuring a thorough exploration of potential siRNA candidates. All these features contributed to the superior performance of OligoFormer.

**Fig. 3.**
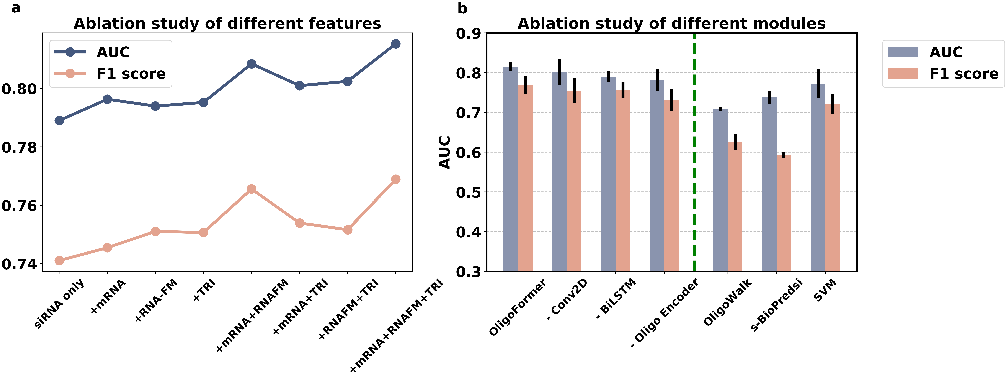
Ablation analysis of different input features and modules of OligoFormer. (a) Ablation study of different features of OligoFormer. The “siRNA only” model utilized only siRNA sequences as the input feature, while the other 7 models incorporated additional features. (b) Ablation study of different modules of OligoFormer. The left side of the dotted green line is the complete OligoFormer and uncomplete OligoFormer with the corresponding module removed. The right side of the dotted green line is previous methods as benchmarks.

Then we evaluated the essentialities of different modules in OligoFormer, including the complete OligoFormer, the model with Conv2D removed, the model with BiLSTM removed, and Oligo encoder removed model with previous methods as benchmarks. As Fig.3b shows, the complete OligoFormer model achieved the best performance, followed by -Conv2D, -BiLSTM, and -Oligo encoder, successively. The model’s performance decreased the most when the Oligo encoder was removed, highlighting its indispensability. The Oligo encoder uses a transformer encoder to capture intricate patterns and dependencies in sequential data, making it well-suited for the complex nature of siRNA sequences.

We also measured how these different features influence the convergency speed of our models. As shown in Figure 1S, adding mRNA features slowed down the convergence speed, while incorporating RNA-FM and thermodynamic parameters accelerated the model’s convergence. The addition of mRNAs appears to introduce complexities that hinder the optimization process, leading to a slower convergence speed. This could be due to the introduction of noise that disrupts the learning dynamics. RNA-FM and thermodynamic parameters likely contribute positively to the convergence speed. As a foundation model, RNA-FM enhances the model’s ability to capture features of RNA sequences and provide valuable representations. And thermodynamic parameters may enable the model to exploit energy-based considerations, guiding the optimization process more efficiently.

### 3.4 Model interpretation

Understanding the inner workings of models is crucial for gaining insights into the decision-making process. We visualized the interpretability of OligoFormer by generating the saliency map through gradient backpropagation on Huesken dataset. Saliency map serve as a valuable tool to elucidate which parts of the input siRNAs significantly influence predictions of OligoFormer and trace the impact of each nucleotide feature back to the model’s input. Regions with higher saliency values indicate molecular components that play a substantial role in the model’s decision-making process. As Fig.4 shows, A and G bases in the first 2 positions of siRNAs showed high saliency value, while the U base in the last 2 positions of siRNAs showed high saliency value. This phenomenon has been reported by previous researches[32], which indicated that A and G in 1-2 position of siRNAs and U in 18-19 position of siRNAs showed a high pearson correlation coefficient(PCC) with the efficacy.

**Fig. 4.**
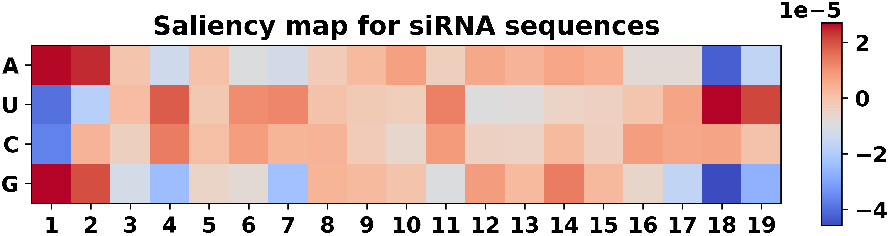
Saliency map for siRNAs on Huesken dataset. Red represents high gradients and importance in the corresponding position.

### 3.5 OligoFormer with off-target searching for siRNA design

We also incorporated 2 existing off-target tools into OligoFormer to achieve better siRNA design. As Fig.5 shows, this pipeline combines OligoFormer with off-target searching to provide an accurate prediction of siRNA efficacy and off-target effects.

**Fig. 5.**
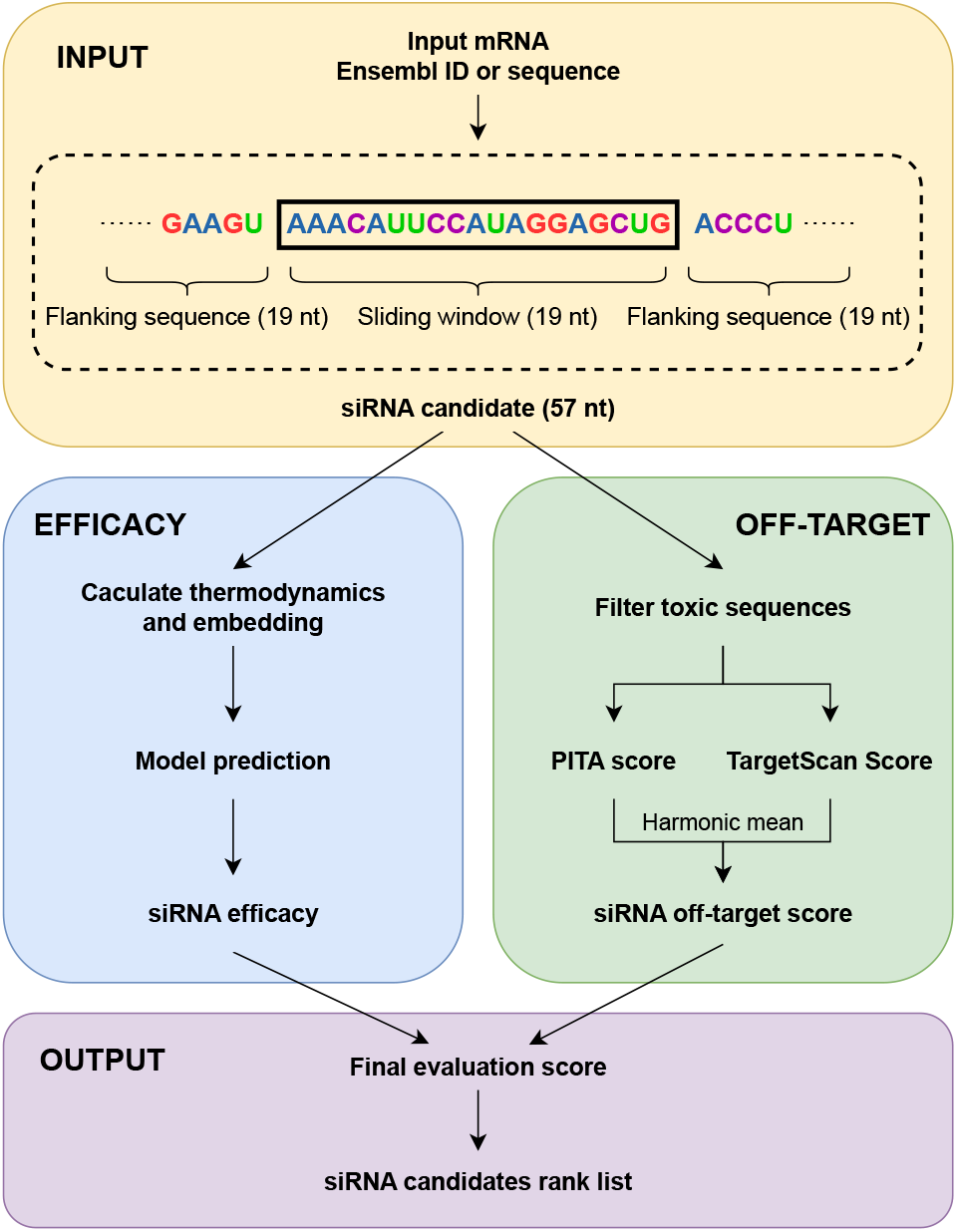
The pipeline of OligoFormer with off-target searching for siRNA design. Given a target mRNA as the input, siRNA efficacy and siRNA off-target score will be calculated to get a final evaluation score and a ranked list of siRNA candidates.

OligoFormer can efficiently infer effective siRNAs given a target mRNA as the input. During the inference process, a sliding window of 19 nucleotides will be utilized to scan the whole target mRNA sequence and generate a set of potential siRNA candidates. Meantime, each siRNA candidate is expanded to its corresponding 57-nucleotide mRNA sequence and the corresponding thermodynamic parameters of these siRNA candidates will be calculated. Then OligoFormer incorporates RNA-FM embedding vectors as pre-trained representations of siRNA sequences. Next, OligoFormer will feed these diverse features into the pre-trained OligoFormer neural network to predict the efficacy of siRNA candidates. The output of the model represents a prediction score for each siRNA, reflecting its potential effectiveness in inducing gene knockdown.

The off-target effects associated with each siRNA are also incorporated into the pipeline to ensure a comprehensive assessment. Any siRNA candidate with motifs which specifically induce immune response is prioritized for filtration. Then some existing miRNA-target prediction methods can be applied to evaluate siRNA miRNA-like off-target potential. TargetScan Context++[33] provides a quantitative model incorporating 14 features for miRNA targeting efficacy prediction, as well as miRNA-like off-target effects, which is the primary module of siRNA off-target effects. PITA[34] is a parameter-free model for miRNA-target interaction prediction based on site accessibility. It considers the difference between the gained free energy from the miRNA-target formation and the energetic cost of opening up the original base pairings of target RNA. Moreover, PITA is a reliable method for siRNA off-target prediction[21]. For each filtered siRNA candidates with a predicted efficacy provided by OligoFormer, two off-target scores are calculated by TargetScan Context++ and PITA respectively within a given interested mRNA set, serving as the basis for customized filtration. After normalization and harmonic average treatment, we finally obtain the off-target scores. Higher output scores from the two methods indicate an increased likelihood of siRNA candidates targeting unexpected mRNAs.

The final output is a ranked list of siRNAs based on a final evaluation score that integrates both the predicted efficacy and the potential off-target risks. The formula for final evaluation score is:

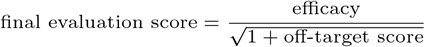

This ranked list provides researchers and practitioners with a prioritized set of siRNA candidates, facilitating the selection of optimal candidates for further experimental validation and therapeutic development.

## 4 DISCUSSIONS

Compared with previous methods, OligoFormer comprehensively considers the RNA sequence features by Oligo encoder, pretrained features by RNA-FM, and thermodynamic parameters. We believe there are three main reasons why our model outperforms the others on both intra-dataset and inter-dataset validation. Firstly, to the best of our ability, we collected and normalized all published siRNA datasets to account for different dataset characteristics. Secondly, we successfully applied the transformer model to the prediction of siRNA efficacy as the component of Oligo encoder, which will leverage the power of transformers to improve the efficiency and accuracy of siRNA design in various applications. And the Oligo encoder contributed most to the model’s performance according to our ablation study. Thirdly, we used RNA-FM to generate pre-trained RNA representations, which also accelerated the convergence speed and model performance. Additionally, we took off-target siRNAs into account and created a comprehensive pipeline for analyzing off-target effects for siRNA design.

However, there is still plenty of room for improvement. The limited representation and quantity of existing datasets severely limit the effectiveness of siRNA design models and further generalization. It is necessary to make more datasets public or build more new datasets. Currently, other pre-trained RNA language models are gaining popularity. RNA-MSM is an unsupervised RNA language model based on multiple sequences that outputs both embedding and attention map[35]. Given that siRNAs and mRNAs are not linked in exactly the same way as non-coding RNA, other models may be able to provide a better characterization of this problem. Chemical modification is an important strategy to reduce the off-target effect of siRNAs, but it has not been included in the prediction process of siRNAs at present. If chemical modification prediction can be encoded and decoded through published data, it may greatly improve prediction of siRNA efficacy and off-target effects, facilitating its application in siRNA design.

This research provides an accurate and robust method based on transformer model for siRNA design. We hold the belief that OligoFormer will provide scientific and comprehensive advice for researchers and help their oligo formed.

## Additional Information

Extended figures and tables are available as supplementary materials.

## Author contributions statement

Zhi John Lu provided patient guidance on this project. Yilan Bai and Haochen Zhong conducted the experiments and wrote the manuscript. Taiwei Wang helped analyse the results.

## Acknowledgments

This work is supported by National Natural Science Foundation of China (32170671, 82371855, 82341101), Tsinghua University Guoqiang Institute Grant (2021GQG1020), Tsinghua University Initiative Scientific Research Program of Precision Medicine (2022ZLA003), Bioinformatics Platform of National Center for Protein Sciences (Beijing) (2021-NCPSB-005). This study was also supported by Bayer Micro-funding, Bio-Computing Platform of Tsinghua University Branch of China National Center for Protein Sciences.

