## Supplementary Data for "OligoFormer: an accurate and robust prediction method for siRNA design"

**Table 1S. Three datasets of siRNAs**

| Dataset | Number of siRNAs | Number of mRNAs |
|---|---|---|
| Huesken | 2431 | 34 |
| Takayuki | 702 | 1 |
| Mixset | 581 | 40 |

**Table 2S. 24 thermodynamic features of siRNAs**

| Individual feature | Position | PCC |
|---|---|---|
| $\Delta\Delta G^{\circ}_{ends}$ | 1 versus 19 | 0.4665 |
| $\Delta G°$ | 1 | 0.4348 |
| $\Delta H°$ | 1 | 0.4069 |
| $U$ | 1 | 0.3634 |
| $G$ | 1 | -0.3024 |
| $\Delta H°$ | all | 0.2813 |
| $U$ | all | 0.2464 |
| $UU$ | 1 | 0.2415 |
| $G$ | all | -0.2151 |
| $GG$ | 1 | -0.1927 |
| $GC$ | 1 | -0.1988 |
| $GG$ | all | -0.1925 |
| $\Delta G°$ | 2 | 0.1891 |
| $UA$ | all | 0.1759 |
| $U$ | 2 | 0.1761 |
| $C$ | 1 | -0.1783 |
| $CC$ | all | -0.1707 |
| $\Delta G°$ | 18 | -0.1594 |
| $CC$ | 1 | -0.1614 |
| $GC$ | all | -0.1613 |
| $CG$ | 1 | -0.1556 |
| $\Delta G°$ | 13 | 0.1530 |
| $UU$ | all | 0.1508 |
| $A$ | 19 | -0.1282 |

*PCC (Pearson correlation coefficient) for efficacy and each thermodynamic features on the Hu dataset

**Table 3.** Parameter settings of model and training

| | Module | Parameter |
|---|---|---|
| **Model parameters** | 2D Convolution Layer | Kernel size = 2x5, Stride = 1, Filters = 32, Pad = 0 |
| | Max Pooling Layer | Pool size = 2x2 |
| | Average Pooling Layer | Pool size = 2x2 |
| | Bidirectional LSTM | Hidden dimension = 32 |
| | Multi-head Transformer Encoder | Layers: 3 layers, Hidden dimension = 64 |
| **Training parameters** | Optimizer | Adam |
| | Loss function | MSE |
| | Learning Rate | 0.0001 |
| | Weight decay | 0.999 |
| | Batch Size | 16 |
| | Epochs | 200 |
| | Early stop | 30 |
| | Seed | 42 |

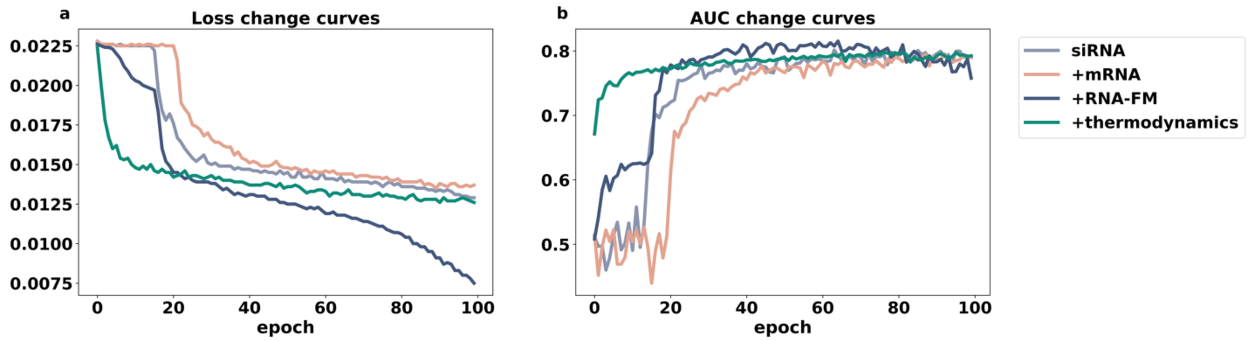


**Figure 1S. Model convergence speed curve.** (a) The loss change curves of models with different features. (b) The AUC change curves of models with different features. The addition of mRNA features will slow down the convergence speed of the model, and the addition of RNA-FM and thermodynamic features will greatly accelerate the convergence of the model.

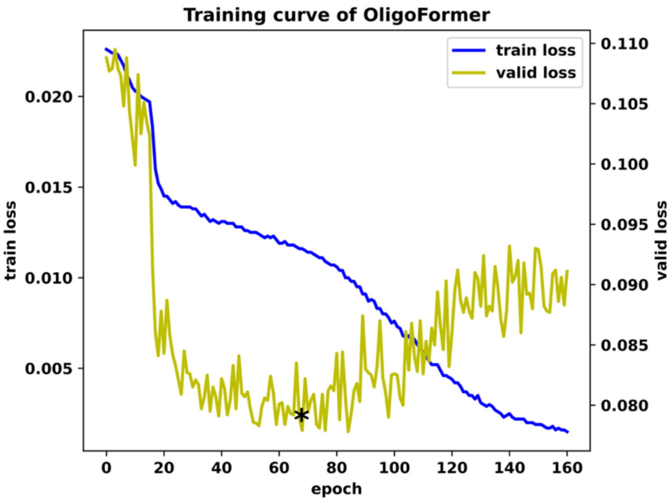


**Figure 2S. The training curve of OligoFormer.** The blue line and yellow line represent training loss change validation loss change during training, respectively. The vertical axis on the left represents the value of the training loss, the right vertical axis represents the value of the validation loss, and the asterisk indicates the location where the best model was selected with a loss of 0.793 and an AUC of 0.8153.

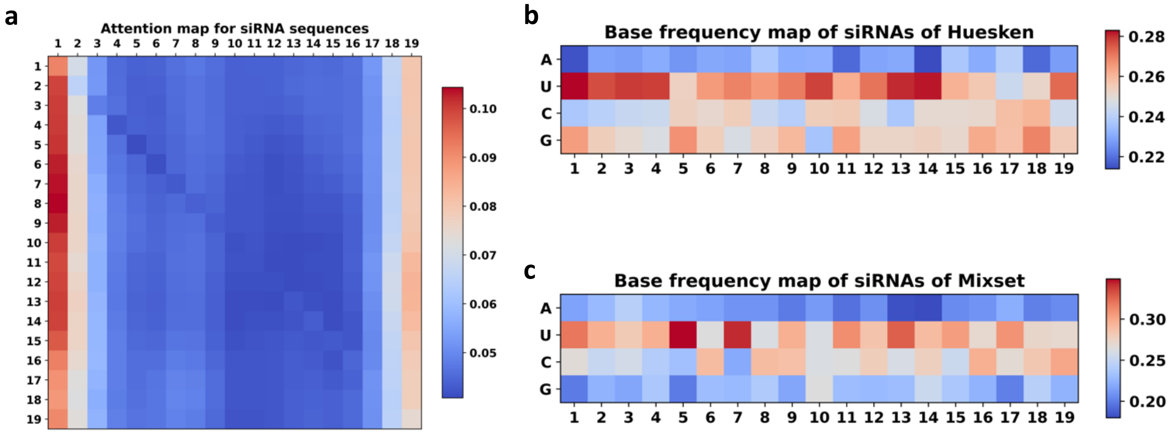


**Figure 3S. Attention map and base frequency map of siRNA sequences.** (a) Attention map of siRNA sequences on Huesken dataset. (b) base frequency map of siRNA sequences on Huesken dataset. (c) (b) base frequency map of siRNA sequences on Mixset dataset.
